# Metagenomes in the borderline ecosystems of the Antarctic cryptoendolithic communities

**DOI:** 10.1101/725200

**Authors:** Claudia Coleine, Davide Albanese, Silvano Onofri, Susannah G. Tringe, Christa Pennacchio, Claudio Donati, Jason E. Stajich, Laura Selbmann

## Abstract

Antarctic cryptoendolithic communities are microbial ecosystems dwelling inside rocks of ice-free areas in Continental Antarctica. In Antarctica, these ecosystems were first described from the McMurdo Dry Valleys, accounted as the best analogous of the Martian environment on Earth and thought to be devoid of life until the discovery of these cryptic life-forms. Our results present the first shotgun metagenomes of Antarctic cryptoendolithic lichen-dominated communities from 18 differently sun-exposed rock samples collected during the XXXI Italian Antarctic Expedition (2015-16), along an altitudinal transect from 834 up to 3100 m a.s.l. Here, we provide the raw data obtained with Illumina Novaseq sequencer, followed by initial functional and taxonomic analysis.

Our results extend understanding of the microbial diversity and biological processes in the Antarctic desert and represent an invaluable resource for the scientific community as a baseline for further studies of this kind to examine the mechanisms and pathways necessary for life to adapt and evolve in the extremes.

## Background & Summary

Antarctic cryptoendolithic communities are microbial ecosystems dwelling inside airspaces of rocks in the super-extreme conditions of the ice-free areas of Continental Antarctica. In Antarctica, they were first described from the McMurdo Dry Valleys, which are considered one of best analogue of the Martian environment on Earth and thought to be devoid of life until the discovery of these cryptic life-forms^1^. The cryptoendolithic ecosystems are highly specialized, adapted to exploit a narrow ecological niche, and represent excellent models to investigate how life can persist at the extreme of aridity, solar radiation and temperature^2^. Studies of these communities can inform the range of parameters where life can still exist, the speculation on the possibility of life beyond Earth, and the astrobiological theory of Lithopanspermia (i.e. transfer of life between planets within meteorites). Despite the rising interest on this topic, only recently molecular and genomic based studies shed light on the structure and diversity of some biological functional groups^3-8^, while our understanding on the response of these microbial communities to environmental stresses is still scant. We applied shotgun metagenomic sequencing, for the first time, to Antarctic cryptoendolithic communities, as a part of the “Metagenomic reconstruction of endolithic communities from Victoria Land, Antarctica” project (PI: Laura Selbmann; co-PI: Jason E. Stajich) supported by the Department of Energy’s Joint Genome Institute (DOE-JGI), to examine biodiversity, ecological function and potential stress response strategies of all community members.

A total of high-quality 3,817,654,184 with a mean of 212,091,899 reads per sample, were generated, resulting in a total of more than 10 million of contigs, with a mean GC content of 58.4±3.2 (mean±SD). A total of 21,647,468 genes were predicted of which 9,264,712 with predicted protein products and 2,943,494 assigned to enzymes. Classification of protein content of the samples into the 24 COGs and 44 KEGG functional categories revealed remarkable homogeneity in gene functional categories among sampled sites at different environmental conditions.

Prokaryotic reads included 21 bacterial phyla, among which Cyanobacteria/Chloroplast (20 up 80%), Actinobacteria (10 up 60%), and Proteobacteria (5 up 35%) were the most abundant, while archaea were represented by Thaumarchaeota alone (∼0.05% of reads).

Eukaryotic reads, based on 18S, were dominated by green algae, Chloroplastida, represented mostly by phylum Chlorophyta and 7 classes, among which Coleochaetophyceae (27-50%) and Trebouxiophyceae (7-40%) were predominant. The ITS dataset indicated Ascomycota as the most abundant fungal group, outweighing other groups such as Basidiomycota which represented a small fraction of the total sequences.

Future studies will be required to examine the mechanisms and pathways necessary for life to adapt and evolve in the extremes and to reconstruct the eukaryotic and prokaryotic genomes from these environmental samples to give important insights on functional roles for the most abundant ones that might perform key ecological processes in these ecosystems.

Predicting how the function, biodiversity, and stress adaptation will respond to different environmental pressures will give insights for predicting and monitoring the effects of global change. Monitoring and predicting on the effects on these border ecosystems, which are tremendously resistant but very susceptible to any perturbations, is critical as they have little to no possibility of recovery once disturbed beyond the narrow range of adaptation^4,6^. Investigations of the pristine Antarctic environment will contribute to understand scenarios for early evolution of life on Earth and may provide insights into the extremes at which life can still persist, informing models of habitability of putative extra-terrestrial environments.

## Methods

### Sampling design

Victoria Land is a region of Continental Antarctica which fronts the western side of the Ross Sea and the Ross Ice Shelf; this land, positioned between the Polar Plateau and the coast and exposed to a wide spectrum of climatic extremes, including temperature and precipitation regimes, covers a latitudinal gradient of 8° from Darwin Glacier (78°00’) to Cape Adare (70°30’S)^9^. Ice-free areas dominate the landscape of Southern Victoria Land and the high-altitude locations of Northern Victoria Land, while low-elevation coastal soils of Northern Victoria Land see considerable marine and biological influence.

Sandstone rocks were collected in Victoria Land along a latitudinal transect ranging from 74°10’44.0”S 162°30’53.0”E (Mt. New Zealand, Northern Victoria Land) to 77°52’28.6”S 160°44’22.6”E (University Valley, Southern Victoria Land) during the XXXI Italian Antarctic Expedition (Dec. 2015-Jan. 2016) by Laura Selbmann.

An altitudinal transect, from 834 to 3100 m a.s.l. was considered. Samples were collected from differently sun-exposed rocky surfaces, to verify the effect of sunlight on settlement and adaptation strategies. Rocks were excised using a geologic hammer and sterile chisel, and samples placed in sterile bags, transported and stored at −20°C in the Culture Collection from Extreme Environments (CCFEE), Mycological Section on the Italian Antarctic National Museum, MNA, at the Tuscia University (Viterbo, Italy).

Metadata are listed in Table 1.

**Table 1.**
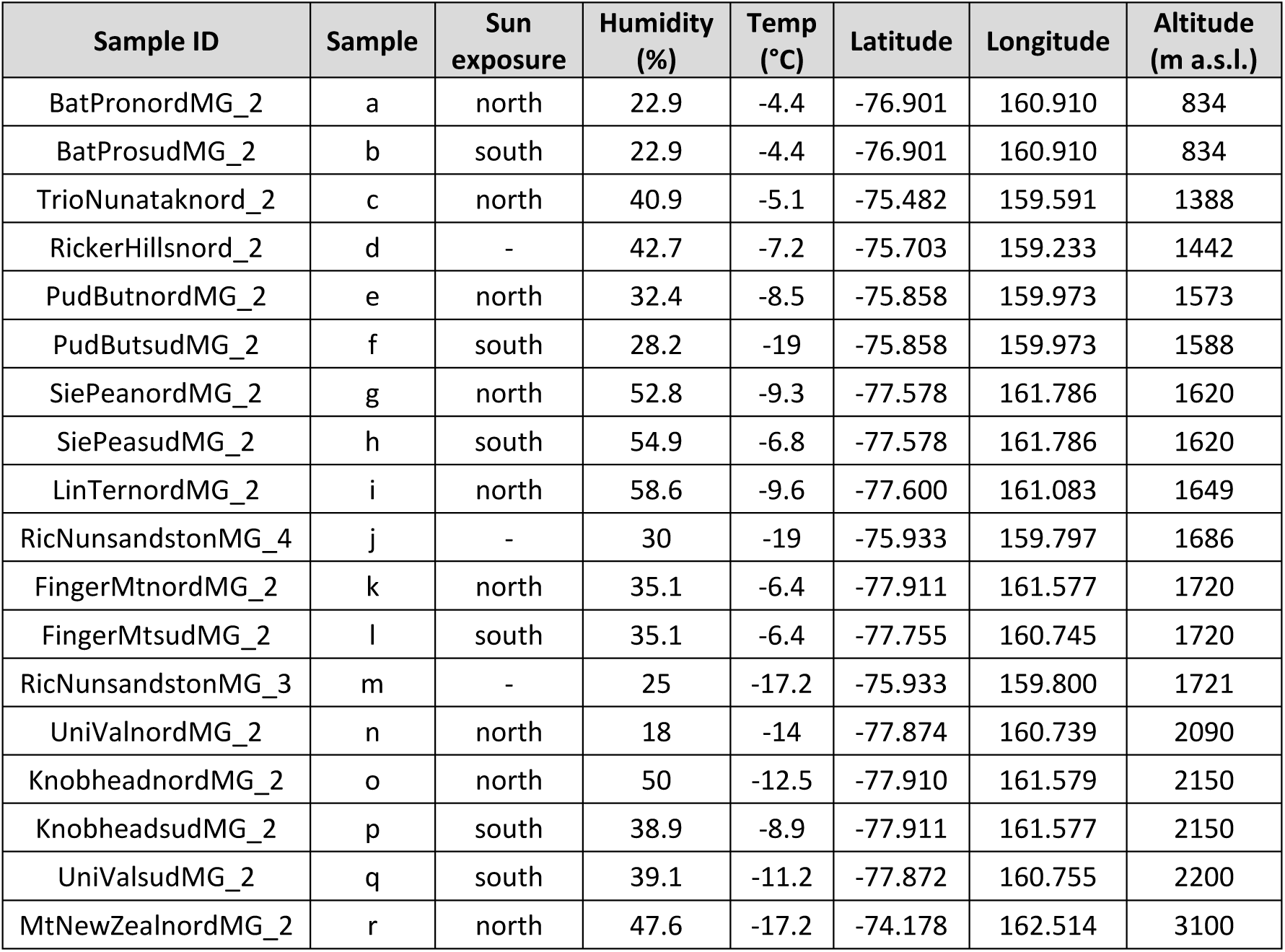
Metadata associated to each sample metagenomes sequenced by Illumina used for this study, including sample sites and environmental conditions.

### DNA extraction, library preparation and metagenomic sequencing

DNA was extracted from three samples for each site and then pooled. We used the MoBio Powersoil kit (MOBIO Laboratories, Carlsbad, CA, USA) and 1 g of carefully crushed rock under sterile conditions with a Grinder MM 400 RETSCH (Verder Scientific, Bologna, Italia) and homogenized before subsampling. The quality of the DNA extracted was determined by electrophoresis using a 1.5 % agarose gel and with a spectrophotometer (VWR International) and quantified using the Qubit dsDNA HS Assay Kit (Life Technologies, USA).

Shotgun metagenomic libraries were prepared and sequenced at the DOE Joint Genome Institute (JGI). Paired-end sequencing libraries were constructed and sequenced as 2×151 bp using the Illumina NovaSeq platform (Illumina Inc, San Diego, CA).

### Downstream analysis

Trimmed, screened, paired-end reads were read corrected using bfc^10^ v.r181 with parameters “-1 -s 10g -k 21 -t 10.” Reads lacking mate pairs after trimming and quality control were also removed. The trimmed, corrected reads in FASTA format were assembled with SPAdes^11^ v3.12.0 with the parameters “-m 2000 -o spades3 --only-assembler -k 33,55,77,99,127 –meta -t 32”. The entire filtered sequence read set was mapped to the final assembly and coverage information calculated using BBMap v38.25 with default parameters and “ambiguous=random”. The version of the processing pipeline is jgi_meta_run.py v2.0.1.

Filtered reads were mapped against 16S, 18S and ITS marker databases using Bowtie2^12^ (v2.3.4.3) (parameters “-k 1 --no-unal”). The 16S, 18S and ITS databases were obtained from the Ribosomal Database Project^13^ (RDP) 16S rRNA gene training dataset v16, the SILVA^14^ eukaryotic 18S subset v123 and the UNITE^15^ database v7.1 respectively and clustered at 95% identity with VSEARCH^16^ v2.10.2. For each marker, mapped (“candidate”) reads were taxonomically classified using SINTAX (parameters “--sintax_cutoff 0.6 --strand both”) against the non-clustered version of the databases. Before classification, paired-end candidate reads were merged using the “fastq join” tool available in VSEARCH and combined with the single-end candidate sequences.

Gene prediction, structural and functional annotation was performed using the DOE-JGI Metagenome Annotation Pipeline^17^ v4.16.5 available into the Integrated Microbial Genomes with Microbiomes^18^ (IMG/M) system v4. Putative amino acid sequences translated from the gene catalog were aligned against: i) the Kyoto Encyclopedia of Genes and Genomes (KEGG) Orthology (KO)^19^; ii) the Clusters of Orthologous Groups (COG)^20^.

## Data Records

Metagenomes sequences used in this study were deposited in GenBank under the accession numbers SRP176584, SRP176586, SRP176587, SRP176592, SRP176590, SRP176595, SRP176596, SRP176601, SRP176600, SRP176598, SRP176606, SRP176608, SRP176612, SRP176609, SRP176611, SRP176667, SRP176669, SRP176664.

IMG annotations are available on JGI IMG/M website (https://img.jgi.doe.gov/) under IMG IDs: 3300030517, 3300032162, 3300031471, 3300031472, 3300031520, 3300031449, 3300031452, 3300031450, 3300033181, 3300031447, 3300030523, 3300031454, 3300031460, 3300031470, 3300031451, 3300031909, 3300031453, 3300031448.

Library size for each metagenome, contigs statistics, taxonomic and functional results can be found in Table S1-S10 (available Online-Only).

All other data that support the findings of this study are available from the corresponding authors upon request.

## Technical Validation

To assess the quality and concentration of genomic DNA, we used both NanoDrop^®^ 1000 spectrophotometer (Thermo Fisher Scientific, Inc.) and Qubit^®^ 2.0 Fluorometer (Invitrogen, Carlsbad, CA, USA). Each sample was quantified in triplicate.

BBDuk v38.25 (“filterk=27, trimk=27”; http://jgi.doe.gov/data-andtools/bb-tools/) was used to remove Illumina adapters, known Illumina artifacts, phiX, contaminants, trim adapters, low-quality sequences and contaminant sequences related to human, dog, cat, and mouse. The procedure discarded reads that contained 4 or more “N” bases, had an average quality score across the read less than 3 or had a minimum length <= 51 bp or 33% of the full read length.

**Figure 1.**
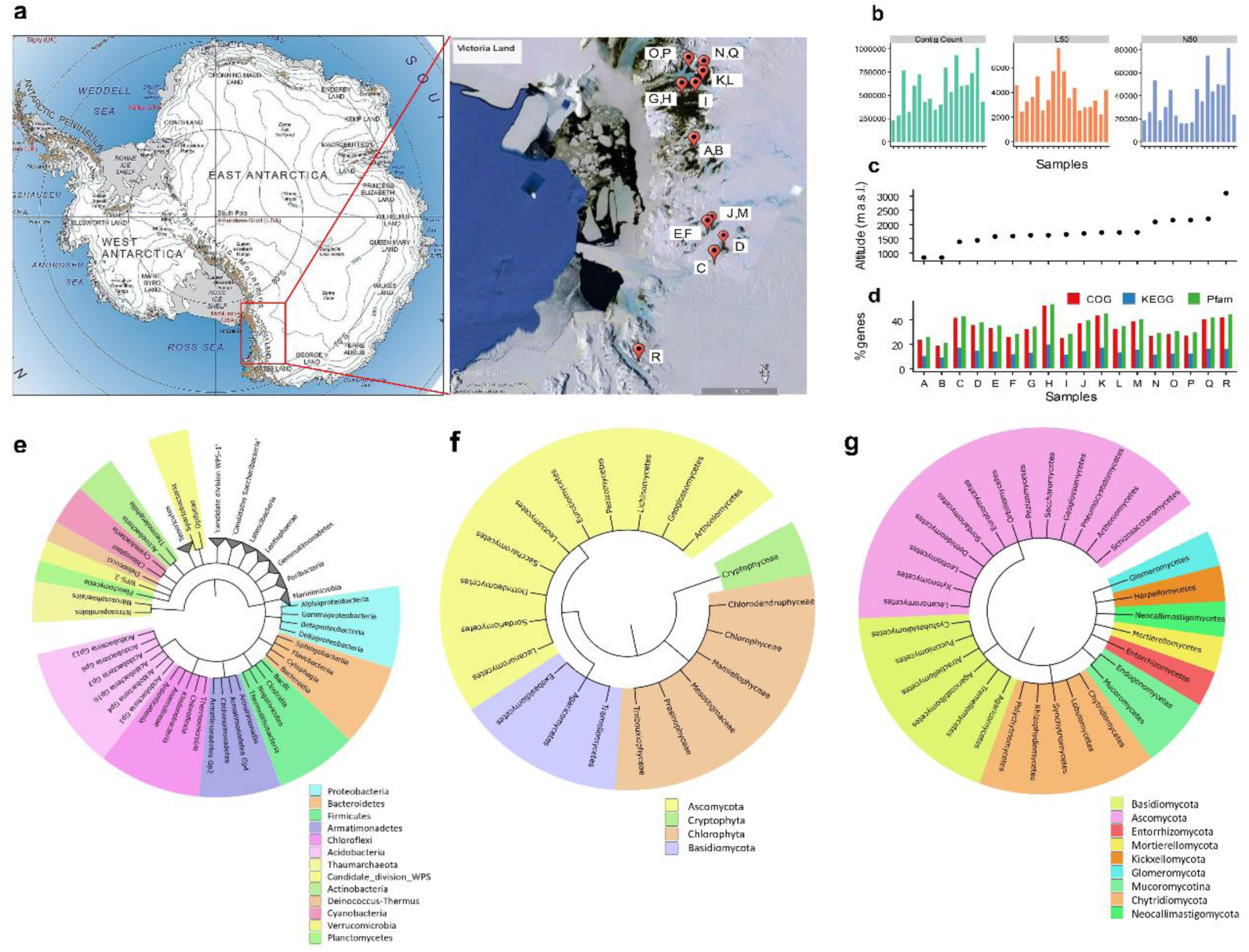
Map of Antarctica and initial findings. **(a)** Map of Antarctica, produced by the LIMA Project (Landsat Image Mosaic of Antarctica) and the sampling area, Victoria Land, Continental Antarctica. Map was generated using Google Earth Pro v7.3 (https://www.google.com/earth/). **(b)** From left to right, number of contigs, the L50 and N50 for each sample. **(c)** Altitudes in the transect considered for sampling. **(d)** Percentages of predicted genes classified in COG, KEGG and Pfam databases. **(e**,**f**,**g)** Cladograms at class level based on 16S, 18S and ITS genomic markers, respectively.

**Figure 2.**
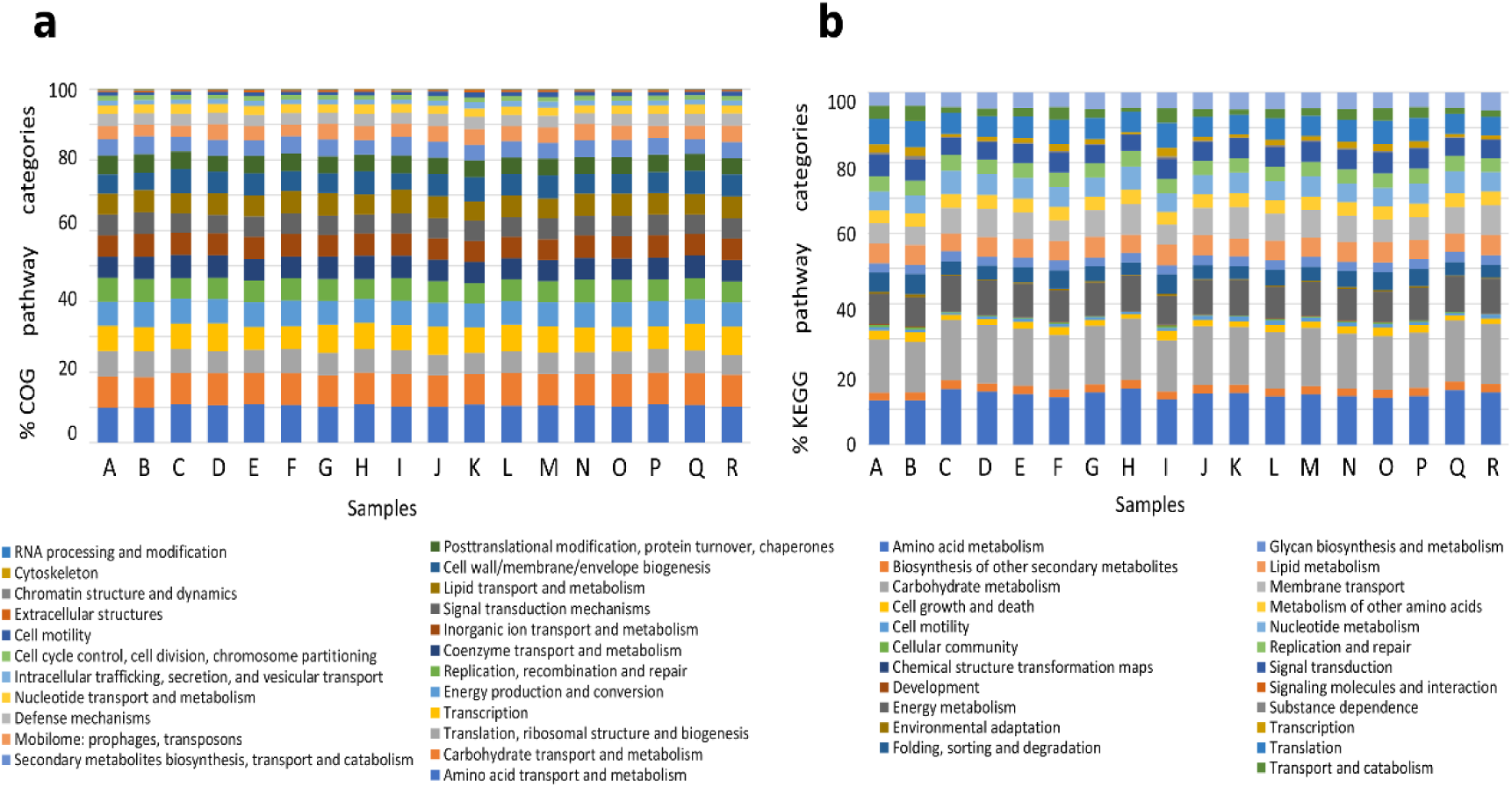
Stacked bar plots representing for each sample the percentage of genes assigned. **(a)** COGs categories. **(b)** KEGG categories.

## Code Availability

No custom code was used to generate or process these data. Software versions used are as follows:

BBDuk v 38.25

SPAdes v 3.12.0

BFC v r181

BBMap v 38.25

jgi_meta_run.py v 2.01

Bowtie v 2.3.4.3

RDP Classifier v16

Silva v123

UNITE v 7.1

VSEARCH v 2.10.2

DOE-JGI Metagenome Annotation Pipeline v4.16.5

IMG/M v4 (https://img.jgi.doe.gov/)

## Acknowledgements

L.S. and C.C. wish to thank the Italian National Program for Antarctic Researches (PNRA) for funding sampling campaigns and researches activities in Italy in the frame of PNRA projects. The Italian Antarctic National Museum (MNA) is kindly acknowledged for financial support to the Mycological Section on the MNA for preserving rock Antarctic samples analysed in this study and stored in the Culture Collection of Fungi from Extreme Environments (CCFEE), University of Tuscia, Italy.

The work conducted by the U.S. Department of Energy Joint Genome Institute, a DOE Office of Science User Facility, is supported by the Office of Science of the U.S. Department of Energy under Contract No. DE-AC02-05CH11231.

Data analyses performed on the High-Performance Computing Cluster at the University of California-Riverside in the Institute of Integrative Genome Biology were supported by NSF DBI-1429826 and NIH S10-OD016290.

## Author contributions

L.S. and J.E.S. are PI and co-PI of the “Metagenomic reconstruction of endolithic communities from Victoria Land, Antarctica” Joint Genome Institute Community Sequencing Project 503708. S.T. and C.P. supported the project as Leader of Metagenomic Group and Project Manager, respectively. Samples were collected by L.S. during the XXXI Italian Antarctic Expedition (2015-2016); L.S., J.E.S and C.C. designed the research; C.C. performed DNA extraction and quality check control; C.C. and D.A. performed data processing and analysis with input from J.E.S. and C.D.; L.S., C.C., J.E.S, and D.A., wrote the paper with input from C.D., S.O., S.T. and C.P.

## Captions Supporting Information (Online-Only)

**Table S1.** Size libraries and sequencing output data from Illumina NovaSeq platform.

**Table S2.** Data generated from metagenomes assembly.

**Table S3.** RDP classification at genus level and relative abundance.

**Table S4.** Silva classification at genus level and relative abundance.

**Table S5.** UNITE classification at genus level and relative abundance.

**Table S6.** List of assigned COGs categories and their relative abundance.

**Table S7.** List of assigned COGs pathways and their relative abundance.

**Table S8.** List of assigned KEGG categories and their relative abundance.

**Table S9.** List of assigned KEGG pathways and their relative abundance.

**Table S10.** List of Pfam functions and their relative abundance.

